# Sex- and Context-dependent Effects of Oxytocin on Social Reward Processing

**DOI:** 10.1101/274027

**Authors:** Xiaole Ma, Weihua Zhao, Ruixue Luo, Feng Zhou, Yayuan Geng, Lei Xu, Zhao Gao, Xiaoxiao Zheng, Benjamin Becker, Keith M Kendrick

## Abstract

We interact socially and form bonds with others because such experiences are rewarding. However, an insecure attachment style or social anxiety can reduce these rewarding effects. The neuropeptide oxytocin (OXT) may facilitate social interactions either by increasing their rewarding experience or by attenuating anxiety, although effects can be sex- and attachment-style dependent. In this study, 64 pairs of same-sex friends completed a social sharing paradigm in a double-blind, placebo-controlled, between-subject design with one friend inside an MRI scanner and the other in a remote behavioral testing room. In this way we could examine whether intranasal-OXT differentially modulated the emotional impact of social sharing and associated neural processing. Additionally, we investigated if OXT effects were modulated by sex and attachment style. Results showed that in women, but not men, OXT increased ratings for sharing stimuli with their friend but not with a stranger, particularly in the friend in the scanner. Corresponding neuroimaging results showed that OXT decreased both amygdala and insula activity as well as their functional connectivity in women when they shared with friends but had the opposite effect in men. On the other hand, OXT did not enhance responses in brain reward circuitry. In the PLC treated group amygdala responses in women when they shared pictures with their friend were positively associated with attachment anxiety and OXT uncoupled this. Our findings demonstrate that OXT facilitates the impact of sharing positive experiences with others in women, but not men, and that this is associated with differential effects on the amygdala and insula and their functional connections. Furthermore, OXT particularly reduced increased amygdala responses during sharing in individuals with higher attachment anxiety. Thus, OXT effects in this context may be due more to reduced anxiety when sharing with a friend than to enhanced social reward.

## Introduction

Humans are highly dependent on their social connections and have a strong motivation to form and maintain close interpersonal relationships such as pair bonds and friendships (Baumeister and Leary, 1995). The affiliative motivation to engage in friendships may be driven both by the positive effects of friendships on well-being (Gangestad and Grebe, 2016) and the hedonic experience of social sharing of emotions (Wagner et al., 2015).

Particularly strong emotional experiences can elicit their being shared socially with close friends, which represents an effective means for helping to communicate as well as regulate them (Rimé, 2009). Numerous previous studies have examined the effects of the presence of a close friend on negative emotional experiences and found that the presence of a companion can attenuate negative affect as well as neuroendocrine and cardiovascular responses to acute stress (Eisenberger, 2013). Initial studies have begun to explore the effects of the presence of a friend during positive and neutral contexts and found that social interactions with peers and sharing positive emotional experiences enhances the hedonic experience and associated neural activity in the brain reward pathways including the ventral striatum, limbic regions and the orbitofrontal cortex (Nawa et al., 2008; Schilbach et al., 2009). A previous neuroimaging study has reported that even awareness that a friend in an adjacent room perceives the same emotional pictures increases positive experience of both negative and positive valence stimuli and activity in reward and emotion regulation-associated brain regions including the ventral striatum and the anterior cingulate cortex (Wagner et al., 2015).

Across species the hypothalamic peptide oxytocin (OXT) facilitates social affiliation and social interactions (Feldman, 2012), possibly via its strong modulatory effects on reward processing (Baracz and Cornish, 2013; Keebaugh and Young, 2011; Rademacher et al., 2015; Scheele et al., 2016; Scheele et al., 2013; Shahrokh et al., 2010) and anxiety (MacDonald and Feifel, 2014; Neumann and Slattery, 2016). Studies administering intranasal OXT in positive and threatening social interactive contexts have reported facilitated processing of positive social cues (Unkelbach et al., 2008), social cooperation (Declerck et al., 2014), increased attention towards positive face expressions (Xu et al., 2015) and recognition of positive and negative facial emotions (Shahrestani et al., 2013), as well as attenuated anxiety (Heinrichs et al., 2003) and neuroendocrine responses to social stressors (Ditzen et al., 2009; Linnen et al., 2011). Moreover, in natural social interaction contexts intranasal OXT has been shown to facilitate the beneficial effects of social support (Heinrichs et al., 2003), positive social communication during couple conflict (Ditzen et al., 2009) as well as social buffering in animals (Smith and Wang, 2014; Smith et al., 2013).

Growing evidence suggests that the specific effects of OXT on human social-emotional behavior develop in a complex interplay with contextual and personal characteristics (Bartz et al., 2011b; Olff et al., 2013). With regard to contextual factors, a number of studies suggest that OXT’s effects on emotional processing depend on the closeness of the relationship with the interaction partner, such that effects were observed for in-group members or close others but not for unfamiliar others. For example, OXT specifically enhances in-group favoritism and parochial cooperation (De Dreu, 2012; Ma et al., 2014) and in-group conformity (Stallen et al., 2012).

In addition, a number of person-related factors can modulate the effects of OXT (Bartz et al., 2011b; Olff et al., 2013), with converging evidence emphasizing the role of sex and attachment style. Recent studies that directly compared the effects of intranasal OXT in men and women reported sex-dependent effects on amygdala activity and connectivity during processing of social information (Gao et al., 2016; Luo et al., 2017). Separate studies have also indicated that OXT may reduce amygdala responses to threatening stimuli in men (Kirsch et al., 2005; Striepens et al., 2012) but increase them in women (Bertsch et al., 2013b; Domes et al., 2010; Lischke et al., 2012). On the other hand, OXT produces sex-independent increased responses in brain reward regions, the nucleus accumbens, putamen or ventral tegmental area, in different studies in men and women in the context of romantic bonds when subjects view the face of their partner (Scheele et al., 2016; Scheele et al., 2013), or when behavior is modified by social reward or punishment (Groppe et al., 2013; Hu et al., 2015). Similar findings of OXT facilitation of nucleus accumbens activation in both sexes in response to social touch by a romantic partner have also been reported in a recent study (Kreuder et al., 2017). It should be noted however that another study has reported sex-differences in the responses of the ventral tegmental area during social reciprocation in the Prisoner’s dilemma game with women, but not men, showing reduced activation (Chen et al., 2017).

Accumulating evidence emphasizes the important role of attachment style as a person-related modulator shaping the specific effects of OXT (Bartz et al., 2011b; Macdonald, 2012). For instance, studies examining endogenous OXT levels found associations with attachment style (Marazziti et al., 2006; Pierrehumbert et al., 2012; Strathearn et al., 2009; Tops et al., 2007), and OXT-administration studies reported pronounced increases in attachment security (Buchheim et al., 2009) and normalization of amygdala hyper-reactivity (Bertsch et al., 2013a; Riem et al., 2016), but also more negative memories about maternal care (Bartz et al., 2010), less positive compassionate experiences (Rockliff et al., 2011) and diminished trust and cooperation (Bartz et al., 2011a) in individuals with an anxious or insecure attachment style.

Taken together, the effects of OXT on social-emotional processing can be mediated both by enhanced striatal reward processing and altered amygdala threat responsivity, with specific effects evolving from a complex interplay between closeness with the interaction partner, sex and attachment style. To disentangle the contribution of these factors the present study employed a randomized placebo controlled between-subject design experiment that combined a single-dose intranasal administration of OXT or placebo (PLC) with a modified version of a previously validated social sharing paradigm (Wagner et al., 2015) during which male and female participants shared their emotional experience either with a close friend or with a stranger.

Given that OXT may particularly strengthen existing affiliative bonds and in-group cooperation and protection we hypothesized that it would increase the emotional impact of sharing experiences with friends rather than strangers. We therefore specifically hypothesized that OXT would promote a positive shift in valence ratings when viewing emotional pictures together with a friend but not with a stranger (compared with viewing alone). Based on previous findings we expected that such an enhanced positive experience in the sharing with a friend context would be associated with either increased reward-related or decreased anxiety-related neural activity. In view of previous studies demonstrating sex-differences in the effects of OXT on anxiety but not reward processing regions we additionally predicted that there would be sex-dependent effects only if it influenced the anxiety circuitry. Finally, given the importance of attachment anxiety for behavior in close relationships and the effects of OXT, we also explored associations with attachment anxiety.

## Materials and Methods

### 2.1 Participants and procedures

Participants were recruited in pairs of good friends (same-sex; duration of friendship > 4 months). A total of 128 pairs of friends (n = 256 participants in total), aged 17-28 years, took part in a randomized placebo-controlled, double-blind, between-subject experiment. All subjects were right-handed and had normal or corrected-to normal vision. Exclusion criteria included current or regular substance or medication use, current or history of medical or psychiatric disorders, any endocrinological abnormalities and MRI contraindications. None of the female participants had been using oral contraceptives or hormone replacement therapy in the 3 months prior to the experiment. All participants provided written informed consent. The study procedures had full approval from the local ethics committee at the University of Electronic Science and Technology of China and were in accordance with the latest revision of the Declaration of Helsinki. The study was preregistered as clinical trial (https://clinicaltrials.gov/ct2/show/NCT03085628; Trial ID: NCT03085628).

In the experiment each pair of friends simultaneously underwent a revised social sharing paradigm (Wagner et al., 2015), with one participant being in the MRI scanner and the other being in an adjacent behavioral testing room. During the paradigm participants were instructed that they would either watch a picture alone, together with their ‘friend’ or together with a ‘stranger’. To provide a context for the ‘stranger’ condition, subjects were told that two pairs of friends (same sex) underwent the experiment simultaneously in different rooms.

In line with recent recommendations (Guastella et al., 2013; Striepens et al., 2011) and the pharmacodynamic profile of intranasal OXT administration (Paloyelis et al., 2016) a single dose of OXT (40 IU, OXT in sodium chloride, glycerin and water, Sichuan Meike Pharmacy Co. Ltd, Sichuan, China) or placebo (PLC, identical nasal spray supplied by the same company but without the neuropeptide) was administered 45 minutes before the start of the experiment. A 40IU dose was chosen since this was the same as in our previous studies showing OXT effects in response to social rejection (Xu et al., 2017) and on processing of social stimuli in the brain salience system (Yao et al., 2018). Although many intranasal OXT studies have used a 24IU dose (Guastella et al., 2013) we have previously found no significant difference between the neural and behavioral effects of 24 vs. 40IU doses (Zhao et al., 2017). During the experimental session each pair of friends received the same treatment. In line with several previous studies (MacDonald et al., 2011), participants could not identify which treatment they received better than chance after the experiment (χ^2^ = 2.176, p = 0.171), confirming successful double-blinding.

To control for potential effects of pre-treatment mood and personal characteristics all participants underwent a comprehensive test battery, including the Positive and Negative Affect Schedule (PANAS; Watson et al., 1988), State-Trait Anxiety Inventory-Trait Anxiety (STAI; Spielberger, 1983), NEO-Five Factor Inventory (NEO-FFI; Costa and McCrae, 1989), Self-Esteem Scale (SES; Rosenberg, 1989), Empathy Quotient (EQ; Baron-Cohen and Wheelwright, 2004), Autism Spectrum Quotient (AQ; Baron-Cohen et al., 2001), Beck Depression Inventory (BDI-II; Beck et al., 1996), University of California at Los Angeles Loneliness Scale (UCLA Loneliness Scale; Russell et al., 1978), and Experiences in Close Relationships Inventory (ECR; Brennan et al., 1998). The McGill Friendship Questionnaire (MFQ; Mendelson and Aboud, 1999) was used both to confirm a high quality of friendship between each pair of participants and to ensure that this did not differ between the two treatment groups. Given that previous studies reported modulatory effects of attachment style on socio-emotional processing (Vrticka et al., 2008) and that individual variations in attachment anxiety have been shown to mediate OXT effects (e.g. Mitchell et al., 2016; Riem et al., 2016), attachment style was assessed using the Adult Attachment Scale (AAS; Collins, 1996; Collins and Read, 1990). To control for potential effects of menstrual cycle and interaction of cycle and treatment, menstrual cycle data was obtained in all female participants. Importantly, no main or interaction effects were observed for menstrual cycle phase or its interaction with treatment **(see details in supplementary results)**.

### 2.2 Stimuli and procedure

A total of 620 pictorial stimuli of comparable quality were selected from the International Affective Picture System (IAPS, Lang, 2005) with additional pictures chosen from the internet to display more Asian protagonists. All pictorial stimuli were rated by an independent sample (n = 34; 18 males) on valence and arousal dimensions (9 - point Likert scale). A total of 72 stimuli per valence condition (positive, negative, neutral) showing people, animals or landscapes were selected. Positive and negative stimuli sets were matched in terms of arousal (positive: mean ± s.e.m. = 6.5 ± 1.79; negative: mean ± s.e.m. = 6.58 ± 1.88, t = 0.734, p = 0.468) and distance from the valence scale midpoint (midpoint = 5; positive: mean ± s.e.m. = 2.16 ± 1.45; negative: mean ± s.e.m. = -2.17 ± 1.56, t = 0.046, p = 0.964). The neutral stimuli were of neutral valence and medium arousal (distance to midpoint valence, mean ± s.e.m. = 0 ± 1.18; arousal, mean ± s.e.m. = 4.72 ± 1.89) **(Table S1)**.

**Table S1.** Stimulus details. Valence ratings, difference to midpoint of valence scale, and arousal ratings. Scales ranging from 1 to 9.

|  | Valence |  | Arousal |
| --- | --- | --- | --- |
|  | mean (s.e.m. ) | Difference to midpoint<br>mean (s.e.m.) | mean (s.e.m.) |
| **Negative Total** | **2.83(0.27)** | **-2.17(0.27)** | **6.58(0.32)** |
| Animal | 2.69(0.28) | -2.31(0.28) | 7.17(0.30) |
| Landscape | 3.29(0.28) | -1.71(0.28) | 5.9(0.35) |
| People | 2.51(0.21) | -2.49(0.21) | 6.66(0.28) |
| **Neutral Total** | **5(0.20)** | **0(0.20)** | **4.72(0.32)** |
| Animal | 4.95(0.26) | -0.05(0.26) | 5.43(0.31) |
| Landscape | 5.04(0.16) | 0.04(0.16) | 3.99(0.33) |
| People | 5(0.18) | 0(0.18) | 4.71(0.28) |
| **Positive Total** | **7.16(0.25)** | **2.16(0.25)** | **6.5(0.31)** |
| Animal | 7.32(0.25) | 2.32(0.25) | 6.82(0.29) |
| Landscape | 6.72(0.26) | 1.72(0.26) | 6.31(0.32) |
| People | 7.44(0.22) | 2.44(0.22) | 6.37(0.30) |
| **Total** | **5(0.39)** | **0(0.39)** | **5.94(0.35)** |

Each trial **(Figure 1)** started with a 2500ms cue indicating the sharing condition (‘ALONE’, ‘TOGETHER with [name of a stranger]’, ‘TOGETHER with [name of the friend]’), followed by the pictorial stimulus displayed for 6000ms. During the stimulus presentation a visual symbol indicating the condition displayed next to the stimulus was displayed as reminder. Following each stimulus participants rated their subjective feelings on a 9 - point scale in terms of valence and arousal (each scale displayed for 3s; valence, 1 = ‘very negative’- 9 = ‘very positive’; arousal, 1 = ‘very low’ - 9 = ‘very high’). Trials were interspersed with a fixation cross (jittered 2500 - 5500; mean 4000 ms) that served as low-level baseline. Stimuli were presented using a mirror-projection system in the scanner room and a personal computer in an adjacent room running E-prime (v2.0, Psychology Software Tools, Inc). To control for confounding effects, names of the unknown ‘stranger’ were matched for length (2 or 3 Chinese characters) with the name of the respective friend and subjects reported in post-scan interviews not knowing a person with the ‘stranger’s’ name. The 216 different stimuli were presented as 6 blocks in a randomized event-related design with each stimulus picture being assigned to one of the three ‘sharing’ conditions (i.e. 6 blocks with 36 pictures in each and each block balanced with respect to stimulus valence). Each block of 36 pictures was additionally balanced for the three categories of stimulus pictures (i.e. people, animals and landscapes). The order of the runs with respect to the stimuli sets used was pseudo-randomized and counter-balanced.

**Figure 1.**
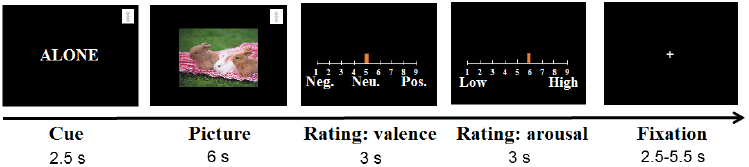
Task procedure. Each trial started with a cue announcing whether the subsequent picture would be watched alone (as in this example) or sharing with the stranger or sharing with the friend. The cues included words in the center of the screen before the picture and a small symbol in the top right corner which was also visible during picture viewing. The picture, whose content could be emotionally positive (as in this example), negative or neutral, was presented, followed by subjective emotion ratings of valence (very negative to very positive) and arousal (very low to very high). The trials were separated by a jittered interval fixation cross in the center.

After the experiment we additionally acquired self-report measures using a modified version of a previous questionnaire to investigate any explicit effects of OXT on sharing with the friend (Jakobs et al., 2001). The scale includes three items on awareness and thoughts about the friend (‘I was thinking of my friend’; ‘I was wondering how my friend would think of the pictures’; and ‘I was wondering how he/she was feeling while seeing the pictures’) and three items relating to intentions to communicate with the friend (‘I wanted to tell my friend what I thought about the pictures’; ‘I had the urge to talk about the pictures with my friend’; and ‘I would like to share the pictures with my friend). For all items subjects provided ratings on a 5-point Likert scale ranging from 1 (not at all) to 5 (always). Ratings collected separately for the three experimental conditions and within them also for the three different valence categories (e.g. How often did you think of your friend when you watched (positive, negative or neutral) pictures together with your friend (or with a stranger or alone)).

### 2.3 Behavioral data analysis

Behavioral data and extracted parameter estimates from the imaging data were analyzed using SPSS 23.0. The primary behavioral variable of interest was the sharing effect, operationalized as relative differences between sharing with a friend or stranger respectively compared to the alone condition (with positive values indicating a shift toward more positive feelings in the sharing condition). The sharing effect was initially evaluated with a one-sample t-test. For the subsequent behavioral analysis data from the subjects in the fMRI scanner and the adjacent room were pooled including group (behavior group vs. fMRI group) as a between-subject factor.

Valence ratings were examined using mixed factor ANOVAs with group (behavior vs. fMRI), sex (male vs. female), and treatment (OXT vs. PLC) as between-subject factors and sharing condition (FRIEND > ALONE vs. STRANGER > ALONE), and valence (positive, neutral, negative) as within-subject factors. Since AROUSAL ratings were significantly decreased by OXT in the ALONE condition the ANOVA analysis included STRANGER, FRIEND and ALONE conditions separately in the condition factor.

Differences in potential confounders were analyzed using independent t-tests. Post-treatment evaluations of awareness and communication motives during social sharing were analyzed using mixed factor ANOVAs with group, treatment and sex as between-subject variables and sharing condition (FRIEND > ALONE vs. STRANGER > ALONE), question category (awareness and communication motives) and valence as within-subject factors. Associations between behavioral and neural indices and attachment style were further examined using bivariate correlation (Pearson for normal, Spearman for non-normal distributed data). The assumption of sphericity was assessed with Mauchly’s test, the Greenhouse-Geisser correction for non-sphericity was applied as necessary, and Bonferroni correction was used for post hoc pairwise comparisons. For significant effects Partial eta-squared (for F test) or Cohen’s d (for t test) was calculated as a measure of effect size. The small, medium, and large effects were represented respectively as 0.01, 0.06, and 0.14 for η^2^, 0.20, 0.50, and 0.80 for Cohen’s d (Cohen, 1988). Tests employed two-tailed p-values, with p < 0.05 considered significant.

### 2.4 fMRI data acquisition and analysis

MRI data was acquired on a 3T MRI system (GE Discovery MR750 scanner, General Electric Medical System, Milwaukee, WI, USA). Functional images were acquired using a T2*-weighted EPI sequence (repetition time, 2, 000 ms; echo time, 30 ms; flip angle, 90°; thickness, 3.4 mm; gap, 0.6 mm; field of view, 240 mm × 240 mm; resolution, 64 × 64; number of slices, 39). To improve normalization of the functional MRI data T1-weighted anatomical images were additionally acquired (repetition time, 6 ms; echo time, 2 ms; flip angle, 9°; thickness, 1 mm; field of view, 256 mm × 256 mm; acquisition matrix, 256 × 256; number of slices, 156).

The images were preprocessed using Data Processing Assistant for Resting-State fMRI (DPARSFA; Chao-Gan and Yu-Feng, 2010; http://rfmri.org/DPARSF). The first six volumes were discarded for signal equilibrium. Preprocessing included slice-time correction, head motion correction using a six-parameter rigid body algorithm, T1-segmentation assisted normalization (resampling at 3 × 3 × 3mm) to Montreal Neurological Institute (MNI) space and spatial smoothing with 8 mm full width at half maximum (FWHM) Gaussian kernel. Participants with head motion exceeding 3.0 mm translation or 3° rotation were excluded.

Statistical analysis was performed using the two-stage mixed-effects General Linear Model (GLM) approach implemented in SPM12 (Statistical Parametric Mapping; Friston et al., 1994; http://www.fil.ion.ucl.ac.uk/spm) implemented in MATLAB 2014a (MathWorks, Inc., USA). The first level matrix included condition-specific onset times (ALONE/STRANGER/FRIEND × positive/negative/neutral) during sharing, separate regressors of no interest for the cue- and rating-presentation times and the movement parameters. In line with the main aim of the study and the behavioral analysis, the second level analysis focused on the main and sex-interaction effects of OXT on sharing-related neural activity.

The sharing paradigm was initially evaluated by examining the main effect of the sharing condition (one-sample t-test including all participants). To determine neural effects of OXT and interactions with sex, context-specific effects (FRIEND > ALONE, STRANGER > ALONE) were examined using full factorial ANOVA models with treatment and sex as between-group factors. To further examine the interaction effect of treatment with sex on functional connectivity between brain regions, we performed a generalized form of context-dependent psychophysiological interaction analyses (gPPI, McLaren et al., 2012, http://brainmap.wisc.edu/PPI). In line with previous studies reporting that OXT affected coupling between the amygdala and insula (e.g. Striepens et al., 2011) and that this effect was mediated by gender (Gao et al., 2016) the analysis of functional connectivity focused on these regions. Significant interaction effects were subsequently disentangled using MarsBar (Brett et al., 2002) to extract individual parameter estimates (beta value) from 8-mm spheres centered at the coordinates of the maximum t value of the corresponding neural effect. The extracted estimates were also used to examine associations with subjective ratings and attachment style.

### 2.5 Thresholding, a priori regions of interest and visualization

All analyses used a whole-brain approach with a significance threshold of p < 0.05 Family-wise error (FWE) corrected for multiple comparisons at a cluster level. To control for false positive rates in the cluster correction approach (Eklund et al., 2016) an initial cluster defining threshold of p < 0.001 was applied to data resampled at 3 × 3 × 3mm (see recommendations by Slotnick, 2017). According to our a priori regional hypothesis, and previous studies reporting consistent effects of OXT on amygdala and insula activity and connectivity (Gao et al., 2016; Wigton et al., 2015), additional analysis focused on the structurally defined amygdala (defined by probabilistic maps as implemented in the Anatomy toolbox (Amunts et al., 2005; Eickhoff et al., 2007; Eickhoff et al., 2005) and insula as well as the striatal-VTA reward system as defined by the Anatomical Automatic Labeling atlas (Tzourio-Mazoyer et al., 2002) (VTA defined according to previous study (Scheele et al., 2013)). For these analyses a small-volume approach (SVC) was used to adjust the FWE-correction (peak level inference, p < 0.05). Connectivity results were visualized with the BrainNet Viewer (BrainNet Viewer: A Network Visualization Tool for Human Brain Connectomics; Xia et al., 2013; http://www.nitrc.org/projects/bnv/).

## Results

### 3.1 Data quality assessment

Eleven subjects were excluded from all further analysis due to missing responses (4 subjects), data acquisition failure (5 subjects), misunderstanding instructions (2 subjects). Given that a key component of the study was that pairs of friends should have a high quality, strong friendship we also excluded 6 pairs of friends with very low mean MFQ scores (average below 120 compared to a mean scores of 200 for other pairs of subjects) from the main analysis, leaving a total of n = 116 in the behavior group (60 males, mean ± s.e.m. age = 21.1 ± 0.2 years), and n = 117 for the fMRI group (61 males, mean ± s.e.m. age = 21.05 ± 0.2 years). Due to excessive head motion during fMRI and data acquisition failure 13 subjects were excluded from the fMRI analysis leaving a total of n = 104 subjects (53 males, mean ± s.e.m. age = 21.0 ± 0.22 years). Excluding these latter subjects from the behavioral analysis had no qualitative effect on the overall results **(see details in supplementary results)**.

### 3.2 Analysis of potential confounders and validation of the paradigm

Treatment groups were comparable in terms of age, mood, personality traits and close relationship characteristics (all ps > 0.078; **Table S2**). Importantly, in the context of the current experiment participants in both treatment groups showed high and equivalent scores on the quality of friendship with the friend with whom they were paired (MFQ scores in behavior group mean±s.e.m.: PLC = 189.36 ± 6.45, OXT = 203.34 ± 7.13, t = -1.455, p = 0.148; MFQ scores in fMRI group: PLC = 200.26 ± 6.70, OXT = 202.07 ± 7.07, t = 0.186, p = 0.853). An analysis of sex differences on questionnaire scores indicated that overall in the behavior groups, women had a higher EQ score (t = -2.081, p = 0.040 Cohen’s d = 0.39) and lower ECR-avoid level (t = 2.789, p = 0.006, Cohen’s d = 0.52) than men.

**Table S2.** Participant characteristics (mean ± s.e.m.)

| Group | Behavior group |  |  |  | fMRI group |  |  |  |
| --- | --- | --- | --- | --- | --- | --- | --- | --- |
| Measurements | PLC<br>(n = 58) | OXT<br>(n = 58) | t | p | PLC<br>(n = 61) | OXT<br>(n = 56) | t | p |
| **Age(years)** | 21.16±0.31 | 21.05±0.26 | 0.257 | 0.789 | 21.02±0.30 | 21.09±0.28 | -0.179 | 0.858 |
| **Positive and Negative Affective Scale (PANAS) –Negative** | 18.19±0.82 | 17.66±0.76 | 0.474 | 0.636 | 18.31±0.77 | 17.11±0.77 | 1.105 | 0.271 |
| **Positive and Negative Affective Scale (PANAS) –Positive** | 27.34±0.70 | 27.84±0.69 | -0.509 | 0.612 | 28.38±0.79 | 27.59±0.68 | 0.751 | 0.454 |
| **State-Trait Anxiety Inventory (STAI)** |  |  |  |  |  |  |  |  |
| State | 37.88±1.01 | 36.55±1.03 | 0.923 | 0.358 | 37.72±0.94 | 38.59±1.04 | -0.622 | 0.535 |
| Trait | 42.45±1.07 | 42.47±1.13 | -0.011 | 0.991 | 41.79±0.92 | 41.63±1.02 | 0.118 | 0.906 |
| **NEO-Five Factor Inventory (NEO-FFI)** |  |  |  |  |  |  |  |  |
| Agreeableness | 42.33±0.53 | 42.90±0.53 | -0.761 | 0.448 | 41.08±0.52 | 41.39±0.54 | -0.412 | 0.681 |
| Conscientiousness | 41.59±0.6 | 40.79±0.68 | 0.872 | 0.385 | 41.44±0.57 | 42.27±0.59 | -1.006 | 0.317 |
| Extraversion | 40.47±0.86 | 40.41±0.80 | 0.044 | 0.965 | 41.21±0.89 | 40.88±0.79 | 0.283 | 0.778 |
| Neuroticism | 32.79±1.09 | 33.47±1.13 | -0.428 | 0.670 | 34.54±0.87 | 33.98±0.93 | 0.440 | 0.661 |
| Openness | 39.28±0.62 | 39.53±0.70 | -0.278 | 0.782 | 40.39±0.68 | 40.05±0.72 | 0.344 | 0.732 |
| **Self-Esteem Scale (SES)** | 30.76±0.61 | 30.19±0.57 | 0.680 | 0.498 | 31.03±0.57 | 31.04±0.60 | -0.004 | 0.997 |
| **Empathy Quotient (EQ)** | 39.78±1.28 | 38.81±1.31 | 0.527 | 0.599 | 38.79±1.35 | 39.45±1.46 | -0.333 | 0.740 |
| **Autism Spectrum Quotient (AQ)** | 19.66±0.76 | 20.29±0.76 | -0.594 | 0.554 | 20.39±0.64 | 19.52±0.70 | 0.924 | 0.358 |
| **Beck Depression Inventory (BDI-II)** | 7.55±0.93 | 8.45±0.90 | -0.691 | 0.491 | 7.33±0.70 | 7.32±0.94 | 0.006 | 0.996 |
| **Adult attachment scale (AAS)** |  |  |  |  |  |  |  |  |
| Anxiety | 2.85±0.10 | 2.92±0.11 | -0.422 | 0.674 | 3.08±0.10 | 2.98±0.11 | 0.670 | 0.504 |
| Attachment & close | 3.41±0.07 | 3.32±0.08 | 0.810 | 0.420 | 3.35±0.07 | 3.19±0.08 | 1.458 | 0.148 |
| **Experiences in Close Relationships Inventory (ECR)** |  |  |  |  |  |  |  |  |
| Anxiety | 3.32±0.12 | 3.23±0.12 | 0.529 | 0.598 | 3.34±0.11 | 3.31±0.14 | 0.135 | 0.893 |
| Avoidance | 3.33±0.11 | 3.05±0.11 | 1.780 | 0.078 | 3.20±0.09 | 3.37±0.11 | -1.185 | 0.238 |
| **McGill Friendship Questionnaires (MFQ)** |  |  |  |  |  |  |  |  |
|  | 189.36±6.45 | 203.34±7.13 | -1.455 | 0.148 | 200.26±6.70 | 202.07±7.07 | 0.186 | 0.853 |
| **UCLA Loneliness Scale** |  |  |  |  |  |  |  |  |
|  | 41.79±1.24 | 42.79±1.16 | -0.589 | 0.557 | 41.66±1.04 | 41.95±2.00 | -0.184 | 0.854 |

### 3.3 Behavioral effects of sharing and their modulation by OXT

We first carried out a preliminary analysis to show OXT had no significant effects compared with PLC on valence ratings in the ALONE condition per se (t = 0.732, p = 0.465). A subsequent ANOVA analysis of valence ratings with group (behavior vs. fMRI) treatment (OXT and PLC) and subject sex as between group factors and picture valence (positive, neutral and negative) and sharing condition (FRIEND > ALONE and STRANGER > ALONE) as within group factors was performed. This revealed a significant main effect of sharing condition [F _(1, 225)_ = 57.811, p < 0.001, η^2^ = 0.204] with ratings being higher when sharing with a friend compared to with a stranger. This confirmed that compared with being alone sharing with a friend produced a stronger increase in valence ratings (i.e. more positive) than sharing with a stranger. There was also a main effect of group [F _(1, 225) =_ 6.437, p = 0.012, η^2^ = 0.028] as a result of sharing effects on valence ratings being stronger in the subjects undergoing fMRI scanning. Examining differences between the PLC groups in the two different testing conditions there was a group × sex interaction [F _(1, 115)_ = 5.594, p = 0.020, η^2^ = 0.046] as a result of valence ratings being significantly higher in the fMRI group than in the behavior group in males (p = 0.004) but not females (p = 0.707).

In terms of treatment related effects, there was a significant group × sex × treatment interaction [F _(2, 251)_ = 12.709, p < 0.001, η^2^ = 0.053] with post-hoc Bonferroni corrected tests indicating that OXT significantly increased sharing effects of valence for females in the fMRI group (t = 9.965, p = 0.002, Cohen’s d = 0.60) and marginally for males in the behavior group (t = 3.977, p = 0.056, Cohen’s d = 0.68) independent of sharing condition and valence. On the other hand, there was no significant OXT effect for females (t = 0.466, p = 0.496) in the behavior group or males (t = 1.847, p = 0.176) in the fMRI group. Additionally, there was a significant sharing condition × treatment × sex interaction [F _(1, 225)_ = 4.649, p = 0.032, η^2^ = 0.020] due to OXT generally increasing the sharing effect with a friend for females (t = 3.880, p = 0.050, Cohen’s d = 0.29) but not with strangers (t = 0.478, p = 0.490), whereas in males there was no significant effect (with stranger: t = 2.335, p = 0.128; with friend: t = 0.155, p = 0.6950). Finally, there was a significant group × sex × treatment × sharing condition × valence interaction [F _(2, 450)_ = 3.024, p = 0.050, η^2^ = 0.013]. Follow-up pairwise Bonferroni corrected post-hoc comparisons showed that in females, OXT only increased valence ratings for positive and neutral valence stimuli in the FRIEND > ALONE sharing condition (behavior group, positive, t = 4.106, p = 0.044, Cohen’s d = 0.82; fMRI group, positive: t = 9.422, p = 0.002, Cohen’s d = 0.65; neutral: t = 5.627, p = 0.031 Cohen’s d = 0.42) as well as in the STRANGER > ALONE sharing condition [fMRI group, positive: t = 4.748, p = 0.030, Cohen’s d = 0.54] **(Figure 2)**. For males the only significant effect of OXT was to increase valence ratings of positive valence stimuli in the STRANGER > ALONE sharing condition in the behavior group (t = 8.323, p = 0.004, Cohen’s d = 0.68). No further main or interaction effects involving OXT were observed.

**Figure 2.**
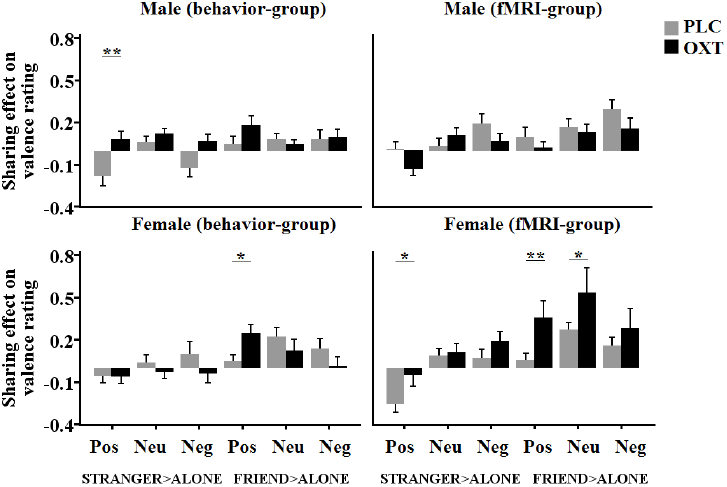
Sharing effect on valence rating. Difference in valence ratings from ALONE condition in male and female subjects in either the behavior or fMRI groups and with either PLC or OXT treatment for positive, neutral and negative stimuli in different sharing-conditions. P = positive valence; NEU = neutral valence and N = negative valence. *p < 0.05, OXT vs. PLC, two-tailed t test. Bars indicate mean ± s.e.m.

An analysis of the arousal ratings differences revealed that OXT significantly reduced ratings in the alone condition compared with PLC (t = 2.183, p = 0.03, Cohen’s d = 0.29). We therefore carried out an ANOVA analysis with condition (ALONE, STRANGER, FRIEND), valence, sex and treatment as factors. This revealed a marginally significant main effect of treatment (F = 3.421, p = 0.066, η^2^ = 0.015) and exploratory post-hoc t-tests confirmed that OXT reduced arousal ratings significantly in the ALONE condition (p = 0.03) and marginally in the FRIEND condition (p = 0.086) but not in the STRANGER condition (p = 0.164) **(Figure 3)**. Thus, OXT appeared to reduce arousal when viewing pictures in the ALONE condition, but without influencing corresponding valence ratings. There were no significant interaction effects involving treatment (see **Supplementary Results** for details on subjective ratings of valence and arousal).

**Figure 3.**
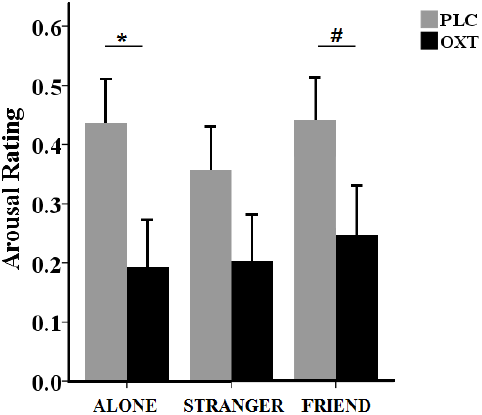
Arousal rating. Arousal ratings in ALONE, STRANGER and FRIEND condition with either PLC or OXT treatment. *p < 0.05 ^#^P<0.1, OXT vs. PLC, two-tailed t test. Bars indicate mean ± s.e.m.

Post-scanning evaluations of social sharing motivation (category: awareness and communication) confirmed stronger motivation when viewing with a friend compared to a stranger [F _(1, 210)_ = 75.883, p < 0.001, η^2^ = 0.265]. There were also main effects of question category [F _(1, 210)_ = 8.365, p < 0.001, η^2^ = 0.038] and valence [F _(2, 420)_ = 8.365, p < 0.001, η^2^ = 0.038]. There was a group × treatment × sharing condition interaction [F _(1, 210)_ = 3.976, p = 0.047, η2 = 0.019] with post-hoc Bonferroni corrected tests showing that OXT generally increased the frequencies of thoughts about friends when sharing with the friend in the behavior group (t = 8.688, p = 0.004, Cohen’s d = 0.55) but not in STRANGER > ALONE sharing condition (t = 0.220, p = 0.639) or in the fMRI group (STRANGER > ALONE sharing condition: t = 0.001, p = 0.981; FRIEND > ALONR sharing condition: t = 0.170, p = 0.681). There was also a group × treatment × category interaction [F _(1, 210)_ = 4.924, p = 0.028, η2 = 0.023] showing that OXT significantly increased the frequencies of thoughts about friends in the behavior group (t = 5.619, p = 0.019, Cohen’s d = 0.48) but not of communicating with them. No effects were observed in the fMRI group (all ps > 0.08). No further main or interaction effects of treatment were observed (all ps > 0.06).

No associations between valence scores and attachment anxiety reached significance for either male or female subjects in either the PLC or OXT groups (ps > 0.09 in all cases). For MFQ scores there was an association with valence scores in females in both PLC (r = 0.293, p = 0.024) and OXT (r = 0.269, p = 0.051) groups although not in males (ps > 0.156).

### 3.4 Functional MRI changes during sharing and the effects of OXT

In line with a previous study (Wagner et al., 2015), the whole brain analysis indicated that across groups sharing with a friend [FRIEND > ALONE] activated left and right medial prefrontal regions extending to the orbitofrontal cortex [peak voxel coordinates, (-18, 57, 6), T = 4.60, cluster FWE corrected p = 0.024, df = 103, k = 197]. In addition, significantly increased activity occurred in the left and right precuneus [peak voxel coordinates, (-6, -60, 33), T = 5.10, cluster FWE corrected p = 0.004, df = 103, k = 332]. For the STRANGER > ALONE sharing condition the left and right precuneus [peak voxel coordinates, (-45, -75, 6), T = 8.13, cluster FWE corrected p < 0.001, df = 103, k = 4356] also showed greater activation.

Examining context-specific effects of OXT revealed no significant effects for STRANGER > ALONE at the whole brain level, whereas for FRIEND > ALONE there was a treatment × sex interaction in the right postcentral gyrus [peak voxel coordinates, (54, -15, 39), T = 4.95, cluster FWE corrected p = 0.004, k = 347] and the right insula [peak voxel coordinates, (39, -18, 12), T = 4.32, cluster FWE corrected p = 0.003, k = 351] **(Figure 4A)**. Extraction of parameter estimates revealed that for FRIEND > ALONE OXT significantly increased right postcentral (p = 0.001) and right insula (p = 0.008) activity in males whereas it decreased activity in females (right postcentral: p = 0.001; right insula: p = 0.005) **(Figure 4B, Figure 4C)**.

**Figure 4.**
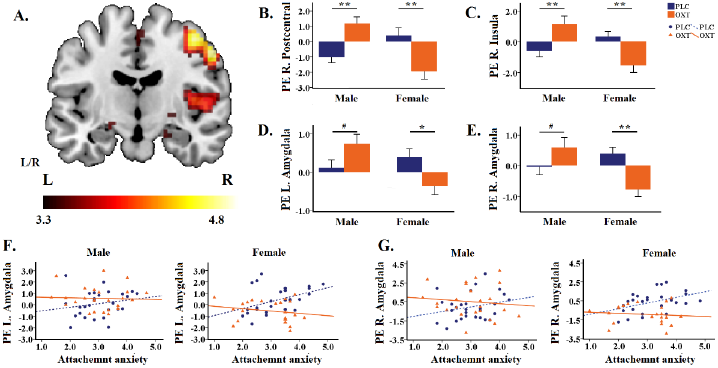
Interaction effect of treatment with sex in brain activation in FRIEND > ALONE sharing condition (A). The t map of the interaction effect showed an activated cluster peaking at the right post central, right insula, right thalamus, left amygdala and right amygdala. Parameter estimates were extracted using an 8-mm-radius sphere centered at the peak MNI coordinates. (B-E) Extraction of parameter estimates revealed significantly changes in (B) right postcentral gyrus (C) right insula (D) left amygdala (E) right amygdala indication that OXT reduced activation in females but not in males (F-G) scatter-plot of a correlation analysis between the anxiety attachment and (F) left amygdala or (G) right amygdala activation in FRIEND > ALONE sharing condition. *P < 0.05, **P < 0.01, ^#^P<0.1, OXT vs. PLC, two-tailed t test. Bars indicate mean ± s.e.m.

Most importantly, for the FRIEND > ALONE sharing condition an amygdala focused analysis revealed an interaction effect in the bilateral amygdala [left amygdala, peak voxel coordinates, (-21, -9, -12), T = 3.61, FWE corrected p_svc_ = 0.0036, df = 100, k = 2; right amygdala, peak voxel coordinates, (21, -9, -15), T = 3.39, FWE corrected p_svc_ = 0.036, df = 100, k = 2] **(Figure 4A)**. Probabilistic mapping on the amygdala sub-regional level indicated that the effect was localized in the centromedial region of the amygdala. Extraction of parameter estimates demonstrated that OXT attenuated bilateral amygdala activity in females (left, p = 0.023; right, p = 0.003), whereas it tended to increase it during sharing with a friend in males (left p = 0.052; right p = 0.097) **(Figure 4D, Figure 4E)**. Since no significant main effects or interactions involving treatment were found for striatal-VTA reward system, no further analysis was focused on them.

Analysis of associations with adult attachment style demonstrated that OXT decreased the positive correlation between attachment anxiety and amygdala activity during sharing with a friend in females (left amygdala: OXT: r = -0.212, p = 0.319; PLC: r = 0.442, p = 0.021; Fisher’s z = 2.309, p = 0.021; right amygdala: OXT: r = -0.134, p = 0.533; PLC: r = 0.404, p = 0.037; Fisher’s z = 1.885, p = 0.059) but not males (left amygdala: OXT: r = 0.045, p = 0.832; PLC: r = 0.257, p = 0.187; right amygdala: OXT: r = -0.092, p = 0.662; PLC: r = 0.233, p = 0.233) **(Figure 4F, Figure 4G).** This suggests that OXT particularly reduced amygdala activation during sharing with a friend in females with higher attachment anxiety scores. Associations between amygdala activity and behavioral valence and arousal ratings or MFQ scores in different sex and treatment groups failed to reach statistical significance.

### 3.5 Effects of OXT on amygdala and insula functional connectivity

During sharing with a friend [FRIEND > ALONE] a significant treatment × sex interaction was found for right insula coupling with the left amygdala [peak voxel coordinates, (-24, -9, -15), T = 3.36, FWE corrected p_svc_ = 0.025, df = 100, k = 4] **(Figure 5A)**. Extraction of parameter estimates demonstrated that OXT increased coupling in males (p = 0.047), whereas it decreased it in females (p = 0.034) **(Figure 5B)**. For the STRANGER > ALONE sharing condition, whole-brain analysis did not reveal any main or interaction effects with the insula as seed regions. Associations between functional connectivity with attachment style and sharing experience in the terms of valence and arousal failed to reach significance (p > 0.086 in all cases).

**Figure 5.**
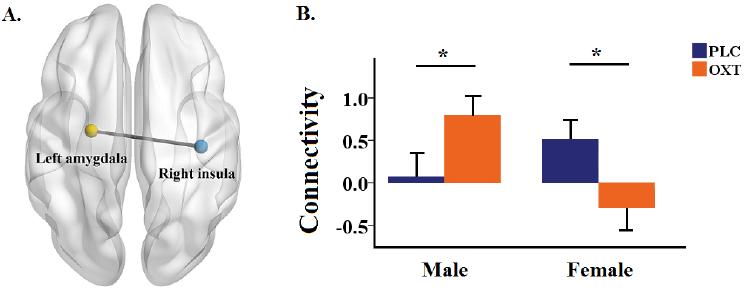
Functional connectivity analysis of treatment × sex interaction effects for FRIEND > ALONE (A) The treatment × sex interaction with right insula and left amygdala as seed regions. (B) Bar graphs illustrate the extraction of parameter estimates for right insula-left amygdala coupling. *p < 0.05, **p < 0.01, OXT vs. PLC, two-tailed t test. Bars indicate mean ± s.e.m.

## Discussion

In the current study, participants were asked to view and evaluate the emotional impact of positive, negative or neutral valence images in the context of sharing the experience remotely with a friend or a stranger as opposed to alone, and to explore the effect of intranasal OXT in these different social sharing contexts. The results from two samples of participants either tested only on behavioral emotional evaluations or additionally in the context of simultaneous fMRI scanning, demonstrated that OXT facilitated the positive emotional impact of sharing experiences with a friend in female but not male subjects, although effects were strongest for positive valence stimuli and in the fMRI context. These behavioral effects of OXT in females were paralleled by decreased activation in the amygdala and insula and reduced functional connectivity between them, but the opposite pattern in males. The effects of OXT on amygdala activation in females were also strongest in participants with higher attachment anxiety. In line with a previous study (Wagner et al., 2015) sharing with a friend was associated with activity in reward related regions but there were no significant modulatory effects of OXT on reward pathways. Thus, in this social sharing paradigm the effects OXT in enhancing the positive impact of sharing experiences with a friend in females may be due more to it facilitating a reduction of anxiety, akin to social buffering, than to increasing the rewarding effect of social sharing.

Both the behavioral and imaging results demonstrate a sharing effect with close friends, suggesting a psychological and neurobiological mechanism underlying the human tendency to want to share emotional experiences with those with whom we have attachment bonds. Irrespective of stimulus valence, subjects tend to make more positive evaluations under conditions where they share an experience with a friend as opposed to a stranger, which is consistent with several previous studies (Wagner et al., 2015). Our post-experiment assessments revealed that subjects generally experienced a stronger awareness of, and desire to communicate with, their friends when viewing stimuli together with them and this was independent of treatment or sex. This further illustrates the basic human motivation to affiliate with friends. The corresponding imaging results showed sharing stimuli with a friend activated frontal regions, including the orbitofrontal cortex, potentially indicating increased social reward (Wagner et al., 2015; Younger et al., 2010). Similarly, the precuneus also showed increased activation when viewing with a friend compared to alone and this region is particularly associated with self-related thoughts and self-referential processing, (Cavanna and Trimble, 2006; Northoff et al., 2006). This might indicate a neural mechanism whereby sharing with close others also engages self-involvement (Müller-Pinzler et al., 2015; Maresh et al., 2013).

Importantly, the friend sharing effect on increasing positive valence relative to the alone condition was significantly enhanced by OXT in female but not male subjects. Interestingly, in male subjects in the behavior group OXT also showed the same facilitatory tendency when sharing with a stranger which might possibly reflect OXT promoting a ‘tend and befriend’ response (Palgi et al., 2014; Taylor and Master, 2011). Notably, we found corresponding results in neuroimaging measures, indicating that in females OXT showed decreased amygdala and right insula activity and the amygdala-right insula coupling when watching with a friend compared with alone. On the other hand, OXT had the opposite effect in men suggesting a sex-dependent modulatory role of OXT on amygdala responses to sharing context irrespective of valence. This is in agreement with a growing number of studies reporting sex-dependent effects of OXT (Rilling et al., 2014) and with our previous findings both for sex-differences in amygdala responses in a social judgment task (Gao et al., 2016) and sub-liminal face-emotion task (Luo et al., 2017).

Consistent with most studies, our results demonstrated that the effect of OXT on social sharing did not vary with the phase of their menstrual cycle. Earlier studies in healthy women have also not shown different effects of OXT on amygdala reactivity in the luteal (Domes et al., 2010) and follicular phases (Bertsch et al., 2013a), indicating that in healthy women menstrual phase may not strongly influence the direction of OXT administration effects. However, more research is needed in females to disentangle the relationships between menstrual cycle, circulating endogenous sex hormones and OXT (Hecht et al., 2017).

Adult attachment style was associated with amygdala, although not behavioral responses in females in the friend sharing condition. Individuals with higher attachment anxiety scores showed greater activation during PLC treatment and this association was absent under OXT. Thus, OXT reduced amygdala responses most during sharing with a friend in those individuals with higher attachment anxiety, potentially indicating a greater anxiolytic effect. Adult attachment style, is an important trait which can strongly influence social bonds and reactions to social partners and an insecure attachment style has often been shown to modulate the behavioral and neural effects of OXT (Bartz et al., 2011b; Buchheim et al., 2009; Riem et al., 2016). Individuals with an anxious attachment style tend to perceive others as unresponsive or inconsistent, worry about being rejected, and show heightened vigilance to signs of support or hostility (Vrticka et al., 2008). OXT and attachment therefore seem to interact in suppressing subjective anxiety and physiological stress responses and OXT may thus have the potential to improve social interactions with and attachment to familiar people such as friends or psychotherapists in female patients with attachment insecurity, such as in borderline personality disorder (Herpertz and Bertsch, 2015). Female, but not male subjects showed a significant positive association between MFQ friendship scores and the friend sharing condition valence rating in both PLC and OXT groups, although not with neural measures. Thus at the behavioral level the strength of friendship between pairs of females may be more influential on the impact of sharing with friends than in males, although associations were quite weak, possibly due to the fact that all pairs of friends had very high MFQ scores.

To the best of our knowledge the current study is the first to simultaneously investigate the behavioral effects of OXT administration on performance of the same paradigm in different testing environments. The effects of OXT on enhancing the positive valence of sharing with a friend were significantly stronger in female subjects performing inside the scanner. In the behavior group the only effect of OXT which reached significance was for female subjects with positive valence stimuli in the FRIEND > ALONE sharing condition and for male subjects in the STRANGER > ALONE sharing condition for negative valence stimuli. In the fMRI groups the effect of OXT on valence ratings was greater and was significant across the three stimulus valences, although post-hoc tests revealed that only positive and neutral valences were significant individually. On the other hand, the effect of OXT on STRANGER > ALONE in males was absent.

It is unlikely that differences between the characteristics of the friends assigned to the two different experimental groups were responsible for context-dependent effects since in terms of friendship scores (MFQ) and mood, anxiety, depression and other measures taken they were equivalent. An alternative explanation is the impact of the different experimental testing contexts. Exposure to magnetic resonance imaging (MRI) conditions can trigger stress responses and interact with performance and neural correlates of cognitive and emotional processes (Critchley, 2009; Muehlhan et al., 2013). While we did not record any indices of stress in the two contexts only males showed a significant behavioral difference between them in the PLC groups, with sharing stimuli producing greater positive valence effects in the fMRI group. Furthermore, in females both amygdala and insula responses when sharing with a friend in the PLC group were increased and functional connections between them were stronger, which could indicate that they were more anxious than the men. There was also a stronger association between attachment anxiety scores and increased amygdala responses in females than in males. Thus, a possible explanation for the sex-dependent effects of OXT in this context could be that it acts to reduce anxiety more in females compared to PLC treatment by enhancing the effect of social buffering, thereby making sharing with a friend a more positive experience. In males on the other hand the act of sharing by itself may produce a sufficient social buffering effect to reduce anxiety and therefore OXT has no additional effect. Indeed, in males OXT actually significantly increased amygdala and insula responses and thus may have even produced slight anxiogenic effects as has previously been reported (Striepens et al., 2012). It is notable that OXT also reduced arousal ratings independent of valence when subjects were in the ALONE condition where any social buffering would be minimal and so it is possible that OXT treatment alone may have had a slight anxiolytic effect. This is in agreement with a previous study reporting that OXT not only enhanced the anxiolytic effect of social buffering but also produced anxiolytic effects even in the absence of social buffering (Heinrichs et al., 2003).

Reduced amygdala activation in women during sharing with a friend was also accompanied by reduction in insula activity and insula-amygdala coupling after OXT. The insula plays an important role in evaluating tasks with an element of emotional processing (Phan et al., 2002; Singer et al., 2004). A meta-analysis of the whole brain imaging findings following OXT administration found that the left insula is the region most robustly influenced (Wigton et al., 2015). Some studies have reported increased activity in the insula in both women (Domes et al., 2010; Riem et al., 2011) and men (Rilling et al., 2012; Striepens et al., 2012) but others have the results that left insula activity is attenuated after OXT administration in a trust game (Baumgartner et al., 2008) and unreciprocated cooperation (Chen et al., 2016). Furthermore, functional connectivity between the right insula and left amygdala decreased after OXT administration in women when sharing with a friend suggesting that the insula may have been using some of the modulatory functions of the amygdala to bias emotional processing (Striepens et al., 2012).

There are some limitations which should be acknowledged in our current study. First, while our data suggest the hypothesis that in the scanner OXT may have particularly reduced anxiety in females in combination with a social buffering effect, we did not take any physiological measures of stress to support this. Secondly, while we found no menstrual cycle-related effects of OXT we cannot rule out the possibility that hormonal changes may have had some influence of behavioral and neural findings. Thirdly, the sharing paradigm used did not involve friends actually being together with one another and it is possible that this might also have influenced our findings. Finally, our emotional stimuli were only of moderate positive or negative valence to avoid ceiling effects and it is possible that stronger valence stimuli, particularly negative valence ones might also have influenced the effects of OXT.

Taken together, our results provide evidence that in the context of a social sharing paradigm where both social reward and social buffering elements are engaged to influence both behavioral and neural measures, OXT probably enhances the impact of sharing with a friend only in females primarily via an anxiolytic action. Furthermore, the effects of OXT in reducing amygdala responses during sharing with a friend were also associated with attachment anxiety in females. This provides further support for the potential therapeutic use of OXT in reducing anxiety in women with insecure attachment, such as in borderline personality disorder. The sex difference we have observed in both the behavioral and neural effects of OXT may have been contributed to some extent by the increased stress associated with being exposed to an MRI scanner environment and differential effectiveness of the social buffering effect of sharing in the two sexes in reducing stress independent of OXT. A fruitful area for future research may therefore be to determine the extent to which anxiolytic or anxiogenic effects of OXT in both sexes are dependent upon current levels of stress being experienced and the efficacy of social buffering effects per se.

## Contributions

X.L.M. and K.M.K. designed the experiment; X.L.M., R.X.L., X. X.Z. conducted the experiment; X.L.M., B.B., W.H.Z., Y.Y.G. and F.Z. analyzed the data; X.L. M., L.X.L. and Z.G. drafted the manuscript. B.B. and K.M.K. interpreted the results and revised it critically and finalized the manuscript for submission. All authors reviewed the final manuscript.

## Pre-registration

At Clinical-Trials.gov database, Trial ID: **NCT03085628,** https://clinicaltrials.gov/ct2/show/NCT03085628

## Acknowledgements

All authors approved the final version of the manuscript. The authors declare no conflict of interest.

## Supplementary Results Behavioral results

### Effects of menstrual cycle and interaction with treatment

Self-report data indicated that none of the female participants was taking oral contraceptives. To further account for potential effects of cycle phase cycle phase on the experimental day was estimated according to a standardized formula (Garver-Apgar et al., 2008). Examining main and interaction effects of cycle phase using a repeated measures ANOVA with treatment and menstrual cycle as between-subject factor and sharing condition as within-subject factor, revealed no significant effect of cycle (OXT, n = 18, follicular phase; n = 25, luteal phase; n = 7, ovulatory phase; n = 3 missing; PLC, n = 25 women, follicular phase; n = 23, luteal phase; n = 8, ovulatory phase; n = 3 missing; all ps > 0.372), arguing against confounding effects of cycle on the analysis of OXT effects.

### Valence rating results

For valence rating results, in addition to effects related to the sharing condition and OXT, we also found a significant main effect of valence [F _(2, 450)_ = 8.161, P < 0.001, η2 = 0.035]. Follow-up analysis showed that subjects had significantly smaller sharing effect when watching positive stimuli than neutral (p < 0.001) or negative (p = 0.013) stimuli. There were also significant sharing condition × sex [F _(1, 225)_ = 7.540, P = 0.007, η2 = 0.032], valence × group [F _(2, 450) =_ 4.827, P = 0.008, η2 = 0.021], sharing condition × valence [F _(2, 450)_ = 8.221, P < 0.001, η2 = 0.035] and sharing condition × valence × sex [F _(2, 450)_ = 8.933, P < 0.001, η2 = 0.038] interactions.

We additionally confirmed that the behavior results remained stable if we excluded the 13 participants with imaging data problems in the fMRI group. The ANOVA revealed significant group × sex × treatment [F _(1, 212)_ = 11.026, P = 0.001, η^2^ = 0.049], sharing condition × sex × treatment [F _(1, 212)_ = 4.051, P = 0.045, η^2^ = 0.019], group × sex × treatment × sharing condition × valence [F _(2, 424)_ = 3.410, P = 0.034, η^2^ = 0.016] interactions. There was also a significant main effect of sharing condition [F (1, 212) = 59.603, P < 0.001, η2 = 0.219], valence [F _(2, 424)_ = 5.779, P = 0.003, η2 = 0.027] and group [F _(1, 212) =_ 5.187, P = 0.024, η2 = 0.024], as well as significant sharing condition × sex [F _(1, 212)_ = 7.920, P = 0.005, η2 = 0.036], sharing condition × valence [F _(2, 424)_ = 7.192, P = 0.001, η2 = 0.033] and sharing condition × valence × sex [F _(2, 424)_ = 9.531, P < 0.001, η2 = 0.043] interactions.

### Arousal rating results

For arousal rating results, we also found a significant main effect of condition [F(2, 450) = 5.540, P = 0.004, η2 = 0.024], valence [F _(2, 450)_ = 847.971, P < 0.001, η2 = 0.790] as well as condition × group [F _(2, 450)_ = 7.096, P = 0.001, η2 = 0.031], condition × sex [F _(2, 450)_ = 7.127, P = 0.001, η2 = 0.031], valence × group [F _(2, 450)_ = 4.595, P = 0.011, η2 = 0.020], valence × sex [F _(2, 450)_ = 3.492, P = 0.031, η2 = 0.015], valence × group × sex [F _(2, 450)_ = 5.811, P = 0.003, η2 = 0.025], condition × valence [F(4, 900) = 16.474, P < 0.001, η2 = 0.068] interactions.

